# Differential activity patterns in the upper and lower blade of the dentate gyrus

**DOI:** 10.1101/2023.06.07.544044

**Authors:** Felipe Fredes, Laura Masaracchia, Marco Capogna, Diego Vidaurre

## Abstract

The dentate gyrus is considered the first stage in the trisynaptic circuit of the hippocampus. Granule cells of the dentate gyrus fire very sparsely with a low probability of overlapping patterns of active neurons. This characteristic supports the most prevalent view of the dentate gyrus function as that of a pattern separator: to generate dissimilar neuronal representations from overlapping input states that represent similar but not identical environments. However, there are two distinct granule cell blades that have been shown to have different activity patterns: the upper blade and the lower blade. These may support different purposes, but their differential function is not well understood. Here we have recorded calcium imaging data from both the upper and the lower blade of the dentate gyrus from two mice (the upper blade from one, the lower blade from the other), while they perform a simple decision-making task. We found that only the lower blade encodes subjective emotional response while the upper blade preferentially encodes the actual response. Importantly, we found that correlations between cells carried more information about these two behavioural variables than individual firings, suggesting that neuron correlations encode here important information above and beyond individual unit activity.

## Introduction

The dentate gyrus (DG) receives information from the entorhinal cortex(Ruth et al., 1988; Leranth and Hajszan, 2007), supramammillary nucleus (Leranth and Hajszan, 2007), and ventral mossy cells (Fredes et al., 2021) among other areas; and relays, through the mossy fibers, into the hippocampal network via CA3 (Lynch et al., 1973). The most prevalent view of the DG function in this circuit is that of a pattern separator: to generate dissimilar neuronal representations from overlapping input states that represent similar but not identical environments (Leutgeb et al., 2007; Edvard, 2009). Such an operation has been ascribed to the hippocampal DG in species ranging from rodents to humans (Bakker et al., 2008). Importantly, polysensory inputs in the DG are mapped onto a large number of granule cells (GCs) that exhibit extremely sparse firing patterns, resulting in a high probability of non-overlapping output patterns (GoodSmith et al., 2017; Nakazawa, 2017).

Most modern studies on GCs function in rodents have used genetically encoded calcium indicators and imaging, which can provide the spatial resolution required to detect sparse coding and non-overlapping active populations. These studies have focused on the upper blade of the DG due to its better accessibility (Pilz et al., 2016; Hainmueller and Bartos, 2018). However, many studies in the past have shown differential activation of the upper and lower blade of the DG. These differences have been detected by early immediate gene expression, especially c-fos and arc (Chawla et al., 2005; Duffy et al., 2013; Bernstein et al., 2019). It is generally agreed that, for example, there is a higher number of active GCs in the upper blade compared to the lower blade during novel environment exploration. When the animal is in the home cage, both blades have a similar number of active neurons; but this proportion changes after the animal explores a novel environment (Montag-Sallaz et al., 1999; Fredes and Shigemoto, 2021; Fredes et al., 2021). However, this is only indirect evidence, and, to the best of our knowledge, there are no studies directly measuring and comparing activity patterns between the upper and lower blade of the DG.

Here we have recorded calcium imaging data from both the upper and the lower blade of the dentate gyrus from two mice (the upper blade from one, the lower blade from the other), while they perform a simple decision-making task. Animals were presented with a stimulus (light cue) and had to make a response (lick), which could be correct or incorrect depending on where the stimulus was presented (right or left), leading to reward or punishment. We performed a decoding analysis separately on the upper and the lower blade, where, using the neural data, we aimed to predict success versus failure in two different conditions: one where failure just led to a lack of reward (soft punishment), and another where failure also led to a dislikeable airpuff (hard punishment). We also compared the amount of information contained regarding behavioural success in the firing rates vs. firing correlations.

We found that only the lower blade encodes subjective emotional response (because the success-vs-failure could only be predicted during the hard punishment condition), while the upper blade preferentially encodes the actual response (regardless of the type of punishment). Importantly, we found that network features (here based on correlations between cells) were more effective for predicting than individual firings, suggesting that neuron correlations encode here important information above and beyond individual unit activity.

## Results

We implanted a GRIN lens above the dentate gyrus of two animals. From one of them, we obtained images from cells in the upper blade (**Figure 1A**) and from the other animal, we imaged the lower blade of the DG (**Figure 1B**). We obtained activity-associated calcium transients from 25 to 51 cells in the lower blade, and from 16 to 62 cells for the upper blade - not all cells were active during each recording session. Animals were trained to perform a simple decision-making task: the mouse had to lick left or right in response to an appropriate stimulus. The decision could be right or wrong depending on the side of the stimulus (light) presented, and so led to reward or punishment. Punishment came as a lack of reward (soft punishment) in some sessions - hereon regarded as non-airpuff sessions-, and as an air-puff (hard punishment) in other sessions-hereon regarded as air-puff sessions. **Figure 1C** shows three representative traces from three different cells in a given session.

**Figure 1.**
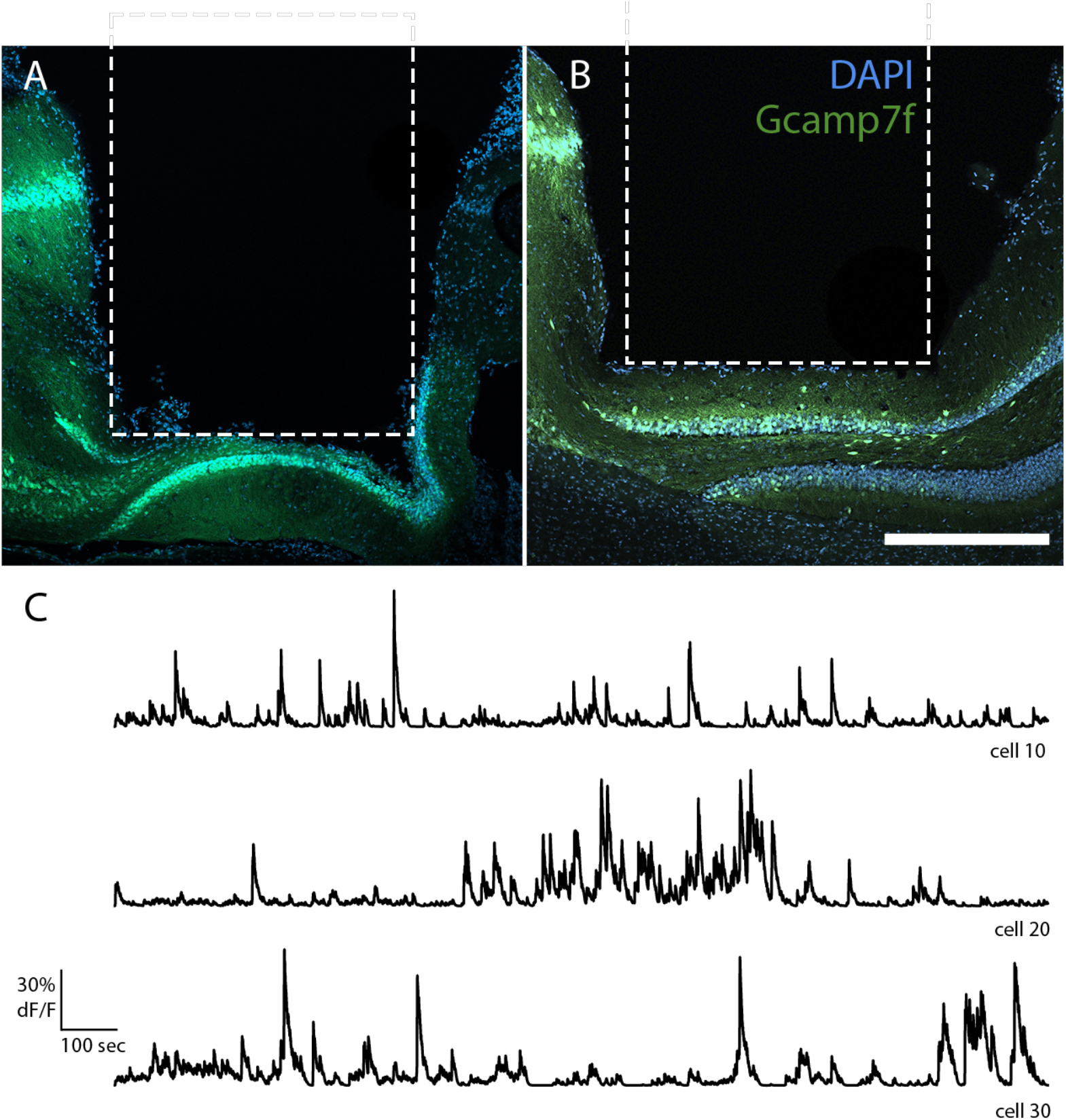
Virus expression, lens position, and representative traces. **A, B**. Confocal image of a coronal section of the dorsal dentate gyrus stained for cell bodies (DAPI, blue) and GCaMP fluorescence (green). The dotted white line depicts the position of the implanted GRIN lens. Scale bar = 500 um. In **A**, the lens is positioned to image the lower blade; in B, it is positioned to image the upper blade. **C**. Shows representative raw traces for three randomly selected cells.

**Figure 2a** shows an example of between-cell correlations in the lower blade during airpuff sessions (upper matrix, during punishment; lower matrix, during response), where correlations were stronger around punishment. To quantify this for each condition and blade, we used PCA analysis, where the explained variance of the first principal component was used as a measure of network integration; see **Methods**. As shown in **Figure 2b**, in the upper blade network integration was significantly higher around response, regardless of the punishment condition (p<0.02). On the contrary, in the lower blade, network integration was significantly higher than the other task-relevant moments in punishment only in the air puff sessions (p<0.02).

**Figure 2:**
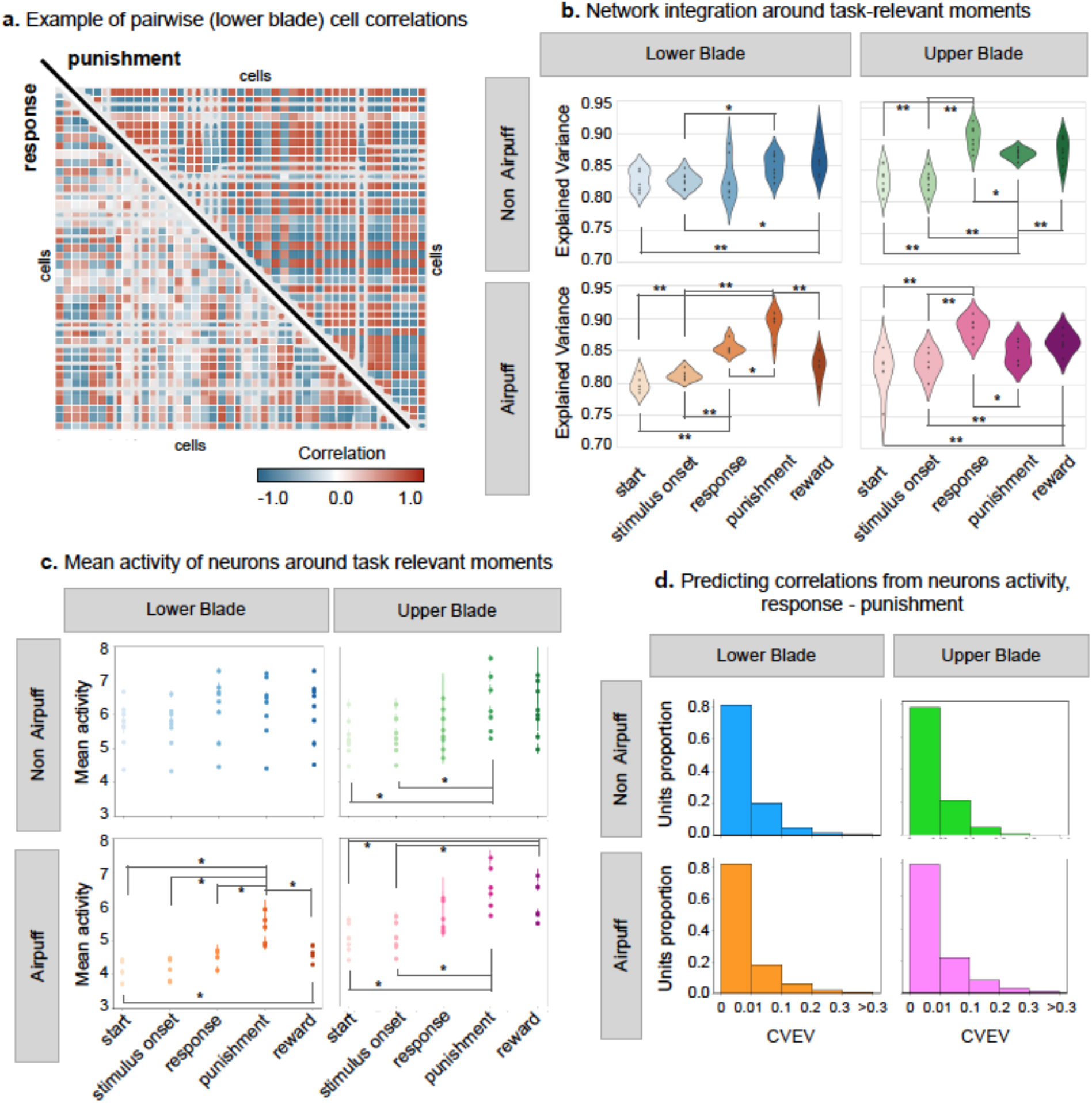
**a**. Example of the average (across trials) cell correlations around the response and punishment in lower blade cells, during a session with air puff punishment (session 14, 41 cells recorded, 115 trials). Correlations are measured in a window of 700 ms, from -500 to +200 ms around response, and -200 to +500 ms around the punishment delivery. **b**. Explained variance of the first principal component of neurons activity (a measure for network integration) around task-relevant moments (windows of 700 ms), averaged across trials per session. Permutation testing was performed at the session level with p values ** <0.01 and * < 0.02. **c**. Average activity of the cells, across trials, and per session, around task-relevant moments (700 ms window). Permutation testing was performed at the session level with p values * < 0.02. **d**. How much variance of the pairwise correlations can be explained by the neurons activity and standard deviation alone. The graph shows Cross Validated Explained Variance (CVEV) of the regression of correlation difference (between response and punishment) on the activity mean and standard deviation difference (between response and punishment).

Average cell activity also had some differences between DG blades and task conditions. As shown in **Figure 2c**, the lower blade cells had similar average activity in all task-relevant moments in the non-airpuff; however, in the air-puff sessions, their activity in punishment was higher (p<0.02). In contrast, the average activity of upper blade cells in both punishment and reward was the highest during the air-puff sessions, while during the non-airpuff sessions, only the activity around punishment was significantly higher than at the start of the trial (p<0.02).

No trivial relation was found between the cells average activity and correlations since mean activity and variance alone were not sufficient to predict pairwise cell correlations: this is shown in **Figure 2d**, where the cross-validated explained variance (CVEV) of the predictions is mostly between 0 and 0.1.

To find if pairwise cell correlations encode distinct information than cell activity, we decoded the binary variable success/failure per trial, either from pairwise correlations (**Figure 3a**) or from cell activity (**Figure 3b**), using a method specifically designed to make the two analyses comparable (see **Methods**). Our decoding results show that in air-puff sessions the decoding accuracy from correlations was considerably higher than from cell activity - for both lower and upper blade cells-, suggesting that more information about the reward or punishment of each trial can be retrieved from correlations then from cell activity itself. For the sake of clarity, we report the difference between decoding accuracy from correlations and from cell activity in **Supplementary Figure 1**.

**Figure 3:**
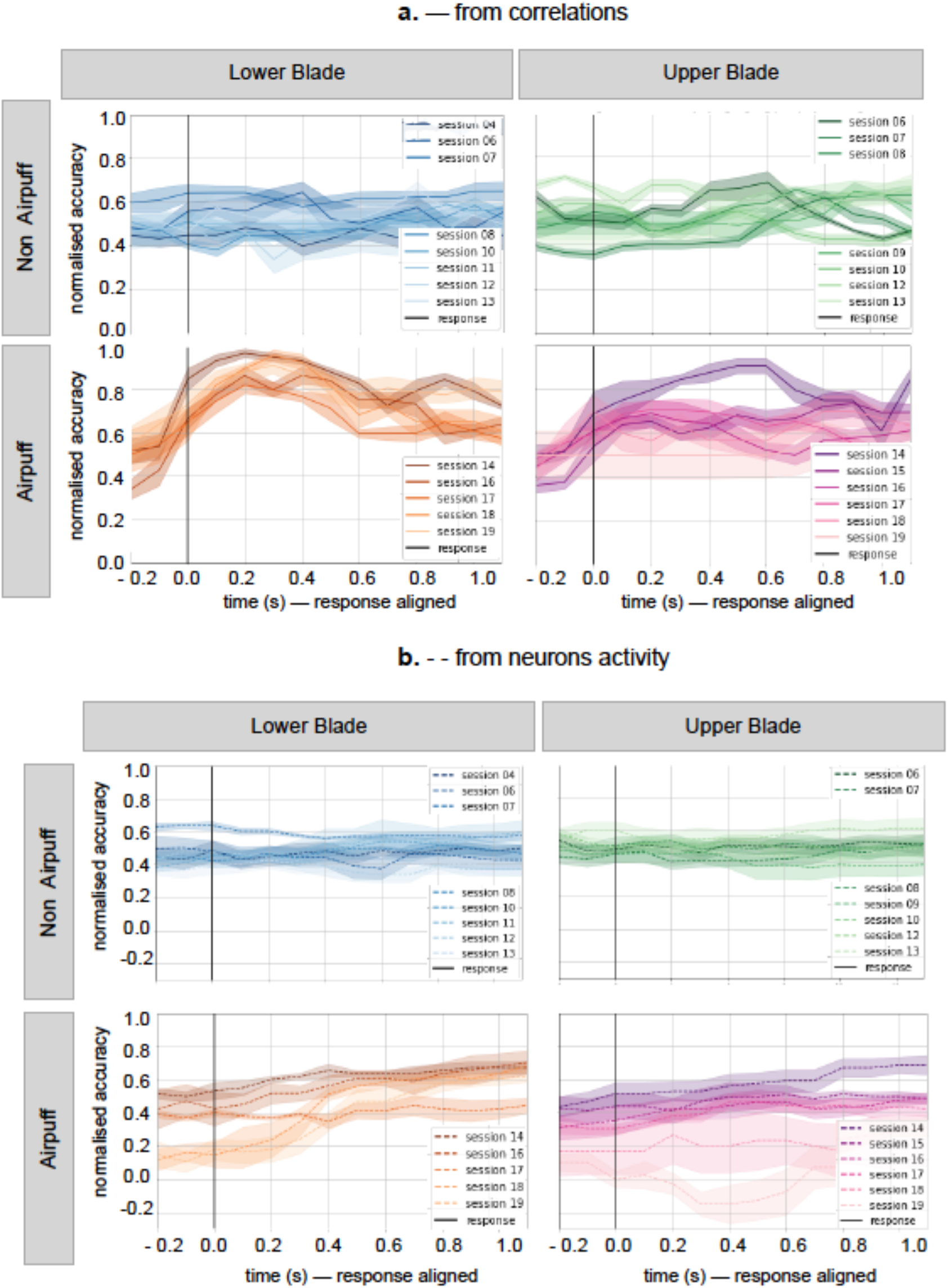
Temporal prediction around response of the binary variable *reward* (vs *punishment*), per each session, using a kernel regression approach, from: **a**. window-based correlation matrices across neurons; and **b**. window-based average activity. The accuracy metric used is *normalised accuracy* = (accuracy - chance level) / (1 - chance level).

## Discussion

Here we have directly compared the activity patterns of the upper and lower blades during a decision-making task, finding: (1) differential predictive information: while the upper blade network activity was more predictive of the actual response, lower blade activity was instead more predictive of the emotional response, i.e. of punishment; (2) stronger between-cell correlations in the lower blade at response time and in the upper blade at punishment time; and (3) that network patterns (here, correlations) contain more information about behavioural success than the individual firings, even though GCs activate very sparsely.

Although there is no published evidence of morphological or hodological differences between the lower and upper blades of the DG, images in the Allen brain atlas of virus injections in the supramammillary nucleus suggest the projections from this structure arborize more densely in the upper blade than in the lower blade. Although the activity of the suprammamilary has been shown to increase with a novel context (Chen et al., 2020), it is difficult to hypothesise a direct function of this synaptic input to the DG. This is because it is not known if the synaptic targets of these terminals are interneurons or GCs, and whether there is a difference in the cells targeted in the lower versus the upper blade.

Another source of inhomogeneity in the DG blades activity can be local circuit differences. We could hypothesise selective microcircuitry connectivity involving specific inhibitory mechanisms that segregate the response of the different blades. For example, it is accepted that local mossy cells help to silence neighbor GCs by activating basket cells interneurons, in a process called “lateral inhibition” (Jinde et al., 2013). If local mossy cells control more inhibitory neurons in the upper blade than in the lower blade, excitatory inputs to the dentate gyrus, carrying emotional responses, could elicit more coactivation of GCs in the lower blade than in the upper one and thus increase correlation for network coding. Detailed high-resolution anatomical studies are needed to unveil these differences.

A robust finding within the existing data is the fact that, in this case, correlations between neurons contained more information about reward than individual firing. We refer here to so-called “noise correlations”, as opposed to “signal-correlations” —which are induced across stimuli, and which information content is mathematically trivial (i.e. if two neurons respond to stimulus A but not to stimulus B, they are signal-correlated)(Nirenberg and Latham, 2003); that is, we show that the pattern of correlation given one condition is different than the pattern of correlation given the other condition, above and beyond the amount of firing. The amount of information carried out by correlations can vary across regions and paradigms, but it is generally acknowledged that the effect of correlations can be substantial in larger populations of neurons (Averbeck et al., 2006); for reviews on understanding and measuring neural correlations see (Cohen and Kohn, 2011) and (Doiron et al., 2016)). Remarkably, here we found a significant effect even for a relatively low amount of recorded cells.

Overall, this paper presents evidence of behaviourally differential activity between the lower and the upper blade of the dentate gyrus. However, we should draw conclusions cautiously since we only have two available animals. We consider this a seed study that will serve as a guide to developing future experiments that can help to explain these unexpected differences in the DG blades functions. For example, are GCs space representations in the upper or lower blades different? Are these inhomogeneities in the response due to microcircuitry differences between the upper and lower blades?

## Methods

### Experimental models

All procedures involving animals were approved by the Danish Animal Experiment Inspectorate. Male mice Prox1-creERT2, developed by Taijia Makinen (Bazigou et al., 2011) were all on a C56BL6/J background and 4 months old for use in this study. Animals received food and water *ad libitum* and were housed under a 12h light-dark cycle. After surgery, animals were housed individually and allowed to recover for at least 3 weeks before experiments.

#### Virus vectors

Retro AAV-hSyn-DIO-GCaMP7f was obtained from Addgene viral service. Viral titration was between 8 × 10^12^ and 7.5 × 10^13^ genome copies per ml.

#### Stereotactic injections, and GRIN lens implantation

Animals were anesthetised using a mixture of 100 mg kg−1 Ketamine / 10 mg kg−1 Xylazine. Animals were placed in a stereotaxic apparatus (Kopf, California) where the skull was exposed and a craniotomy was performed at the following coordinates: -1.8 mm AP and ± 1.3 mm ML. A total volume of 0,5 ul of the virus was injected at a controlled rate of 0.1 µl min−1 using a microinjector (WPI, Sarasota, USA) The pipette was slowly lowered to the target site and remained in place for another 5 min post-injection before being slowly withdrawn. After the virus was injected and the pipette removed, a 1 mm diameter, 4 mm long GRIN lens (Inscopix, California, USA) was implanted. For this, the lens was lowered slowly into the target (1um/sec) and then secured to the skull using Charisma dental cement (Kulzer, Germany). After surgery animals were injected with 5 mg kg−1 of Metacam (meloxicam) subcutaneously as a postoperative anti-inflammatory and painkiller. The animals recovered in a heating pad and then transferred to their home cages and monitored closely for any discomfort signals for the three following days.

### Water restriction and training procedure

Animals were water-restricted until they reached 30% weight loss following a previously published procedure (Guo et al., 2014). Animals were then adapted to the manipulation procedure for 3 days before starting the training protocol. Animals were first accustomed to the experimental settings, being head-fixed and placed in front of the detector, and water rewards were passively delivered to elicit licking. Measurements were then performed in a visual detection task, where the mouse had to actively detect and respond to visual stimuli to obtain rewards at a particular port. Mice learned to lick for rewards at each individual detector, depending on the location of the visual stimulus. For example, if the visual stimulus appeared on the left, the mouse could obtain a water reward by licking at the left lick port during the stimulus-response window (typically 1 s). The first trial after the switch of the stimulus location contained a grace period (∼1.5 s) where licks to either detector during stimulus presentation were rewarded; thereafter, licks to the wrong detector were unrewarded and penalised with a time-out interval or an air puff depending on the trial type. Details of animal preparation and behavioural training can be found elsewhere (Guo et al., 2014).

#### Calcium imaging in behaving mice and preprocessing

Recording sessions began 4 to 6 weeks after lens implantation, right after animals were able to lick on both detector sides. One movie was acquired every day while the animal performed the task for 30 mins, or until the animal was not motivated to lick anymore. Calcium-imaging data were acquired at 20 Hz, constant excitation power, and exposure to compare fluorescent changes across sessions and corrected for motion artifacts using Inscopix Software. FOVs were detected automatically using Inscopix software. Traces were then exported as CSV files for further analysis.

## Analysis

We aimed to compare GCs in the upper and lower blade across different task-relevant moments for two different task conditions (air-puff vs non-airpuff). Selected moments were: trial start, stimulus onset, response, reward, and punishment. For each task-relevant moment, data were considered within a window of 700ms: from 0 to 700 ms after trial start, from -200 to +500 ms around stimulus onset, from -500 to +200ms around response, and from -200 to +500 ms around punishment and reward. Given that active cells were not overlapping across sessions, all analyses were performed at the session level.

### Mean activity and correlation patterns

Average cell activity of the upper and lower blade was compared between task-relevant moments and task conditions. For a given session, the average activity and standard deviation of each individual cell were computed around task-relevant moments. Mean cell activity and standard deviation were calculated within the 700 ms window around task-relevant moments and then averaged across trials. Finally, to assess the difference in overall activity between task-relevant moments, we averaged the mean activity across cells for each session and ran permutation testing. For each session, pairwise correlations of cell activity were computed within the 700 ms window around each task-relevant moment and averaged across trials.

### Predicting correlations from mean activity

In order to rule out that correlation patterns were just a trivial consequence of differences in cell activity rates, we used regression analysis to predict cell correlations from their average activity in two key task-relevant moments (namely, response and punishment). Specifically, for each session, standard ridge regression (cross-validated across trials) was used to predict the difference in pairwise cell correlations between response and punishment from the difference in the average activity and standard deviation (between response and punishment). This yielded a measure of cross-validated explained variance (CVEV) for each pair of cells-averaged across trials-per session.

### PCA analysis

To explore the relevance of cell correlations across task-relevant moments in comparison to individual cell firing, we designed a statistical test based on principal component analysis (PCA). For each task-relevant moment and each trial, we ran PCA on the cell activity —i.e. each PCA run took as input a T x C matrix, where T is the number of time points within the 700ms window and C is the number of cells (time points as samples and cells as features). Then, for each session and task-relevant moment, the variance explained by the first PC was averaged across trials, offering a measure of network integration. Finally, we used permutation testing to assess differences in this measure between each pair of task-relevant moments.

### Decoding from average cell activity and pairwise correlations

Temporal decoding was used to predict the behavioural outcome of each trial (here, success or failure) from both pairwise cell correlations and average cell activity. For each trial within a session, a subset of time points from -200 to +1000 ms around response was considered. Within that subset, we trained and tested a decoder (kernel regression algorithm, see below) per time point. This procedure resulted in a time-point-by-time-point measure of accuracy. Specifically, within the selected -200 to +1000 ms subset around response, we used 700 ms sliding windows to calculate pairwise correlations and average activity. This way, we produced two sets of data: average activity, with dimensions N x C x T (where N = number of trials, C = number of cells and T = number of time points); and pairwise correlations, with dimensions N x C x C x T. In order to reduce correlations dimensionality, we projected pairwise correlations into a Riemann tangent space (Bellaïche, 1996). This transformation provided an alternative representation of the correlation values in a lower dimensional geometrical space while keeping their original relative distances. This way, the Riemann-projected correlations, with only three dimensions (N x R x T, where R =C x (C+1) / 2 is the number of features in Riemann space), were made comparable to the average cell activity, which had the same number of dimensions.

Kernel regression (with a Gaussian kernel)(Shawe-Taylor and Cristianini, 2004) was then used as a decoder, to predict trial success either from the average cell activity or from the Riemann-projected correlations. The algorithm was run in a cross-validated fashion (across trials) for each session. Since the accuracy chance level (i.e., the ratio between number of successful trials and total number of trials) differed for each session, the model performance was not directly comparable across sessions. To address this concern, the accuracy of the model was calculated as the relative increase in accuracy with respect to chance level: relative accuracy = (model accuracy – chance level) / (1-chance level). This way, the accuracy of the model was made comparable across sessions.

---

**Supplementary Figure 1:**
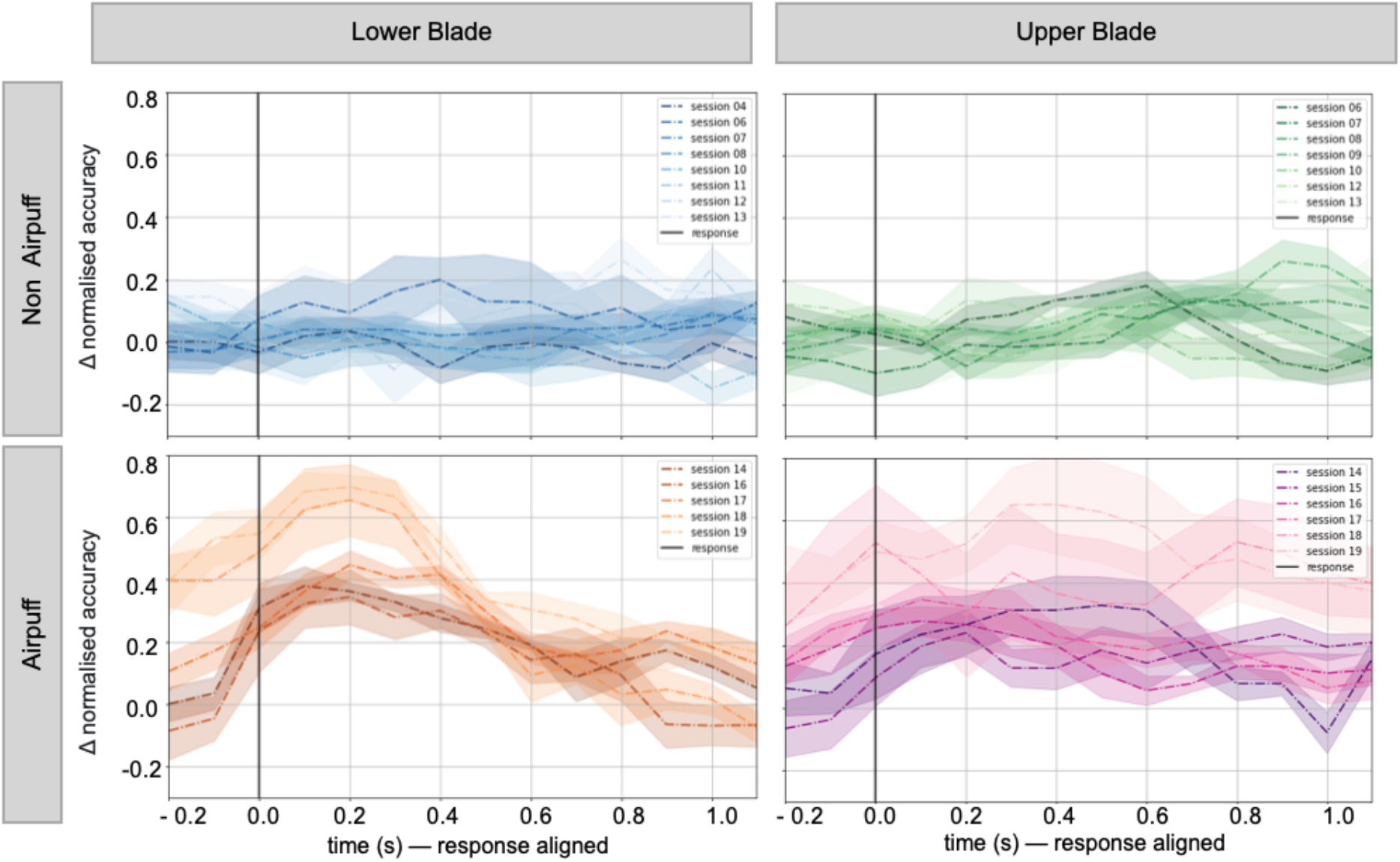
Difference in decoding accuracy between correlations and cells activity, predicting trial reward, per each session and condition. Can be considered an expansion of **Figure 3**, as it shows the difference between plots in **3a** and plots in **3b**.

